# An updated infrageneric classification of the oaks: review of previous taxonomic schemes and synthesis of evolutionary patterns

**DOI:** 10.1101/168146

**Authors:** Thomas Denk, Guido W. Grimm, Paul S. Manos, Min Deng, Andrew Hipp

## Abstract

In this paper, we review major classification schemes proposed for oaks by John Claudius Loudon, Anders Sandøe Ørsted, William Trelease, Otto Karl Anton Schwarz, Aimée Antoinette Camus, Yuri Leonárdovich Menitsky, and Kevin C. Nixon. Classifications of oaks (Fig. 1) have thus far been based entirely on morphological characters. They differed profoundly from each other because each taxonomist gave a different weight to distinguishing characters; often characters that are homoplastic in oaks. With the advent of molecular phylogenetics our view has considerably changed. One of the most profound changes has been the realisation that the traditional split between the East Asian subtropical to tropical subgenus *Cyclobalanopsis* and the subgenus *Quercus* that includes all other oaks is artificial. The traditional concept has been replaced by that of two major clades, each comprising three infrageneric groups: a Palearctic-Indomalayan clade including Group Ilex (Ilex oaks), Group Cerris (Cerris oaks) and Group Cyclobalanopsis (cycle-cup oaks), and a predominantly Nearctic clade including Group Protobalanus (intermediate or golden cup oaks), Group Lobatae (red oaks) and Group Quercus (white oaks, with most species in America and some 30 species in Eurasia). The main morphological feature characterising these phylogenetic lineages is pollen morphology, a character overlooked in traditional classifications. This realisation, along with the now available (molecular-)phylogenetic framework, opens new avenues for biogeographic, ecological and evolutionary studies and a re-appraisal of the fossil record. We provide an overview about recent advances in these fields and outline how the results of these studies contribute to the establishment of a unifying systematic scheme of oaks. Ultimately, we propose an updated classification of *Quercus* recognising two subgenera with eight sections. This classification considers morphological traits, molecular-phylogenetic relationships, and the evolutionary history of one of the most important temperate woody plant genera.

## Part A. History of classifications of oaks

In his original work, Carl von Linné listed 14 species of oaks from Europe and North America: the white oaks *Q. alba, Q. æsculus* (= *Q. petraea* (Matt.) Liebl.), *Q. robur*, and *Q. prinus* (status unresolved); the red oaks *Q. rubra, Q. nigra*, and *Q. phellos*; the Cerris oaks *Q. cerris, Q. ægilops* (= *Q. macrolepis* Kotschy), *Q. suber*; and the Ilex oaks *Q. ilex, Q. coccifera, Q. gramuntia* (= *Q. ilex*), and *Q. smilax* (= *Q. ilex*) (Linné 1753). This number had increased to 150 species when Loudon (1838, 1839) provided the first infrageneric classification of oaks recognising ten sections based on reproductive and leaf characters. Eight of Loudon’s sections (*Albæ, Prinus, Robur*; *Nigræ, Phellos, Rubræ*; *Cerris*; *Ilex*) were based on species described by Linné (**Fig. 1**). New additions were the (fully) evergreen southeastern North American “Live Oaks”, sect. *Virentes*; and the “Woolly-leaved Oaks”, sect. *Lanatæ*, of Nepal (including an Ilex oak and a species that was later recognised as a cycle-cup oak). Loudon’s classification is remarkable in one aspect: he established the fundamental subdivision of European oaks (his sections *Cerris, Ilex*, and *Robur*). This subdivision, although modified, occurs in nearly all later classifications and corresponds to clades in most recent molecular-phylogenetic trees (cerroid, ilicoid, and roburoid oaks; cf. Denk and Grimm 2010; A. Hipp and co-workers, work in progress).

**Figure 1:**
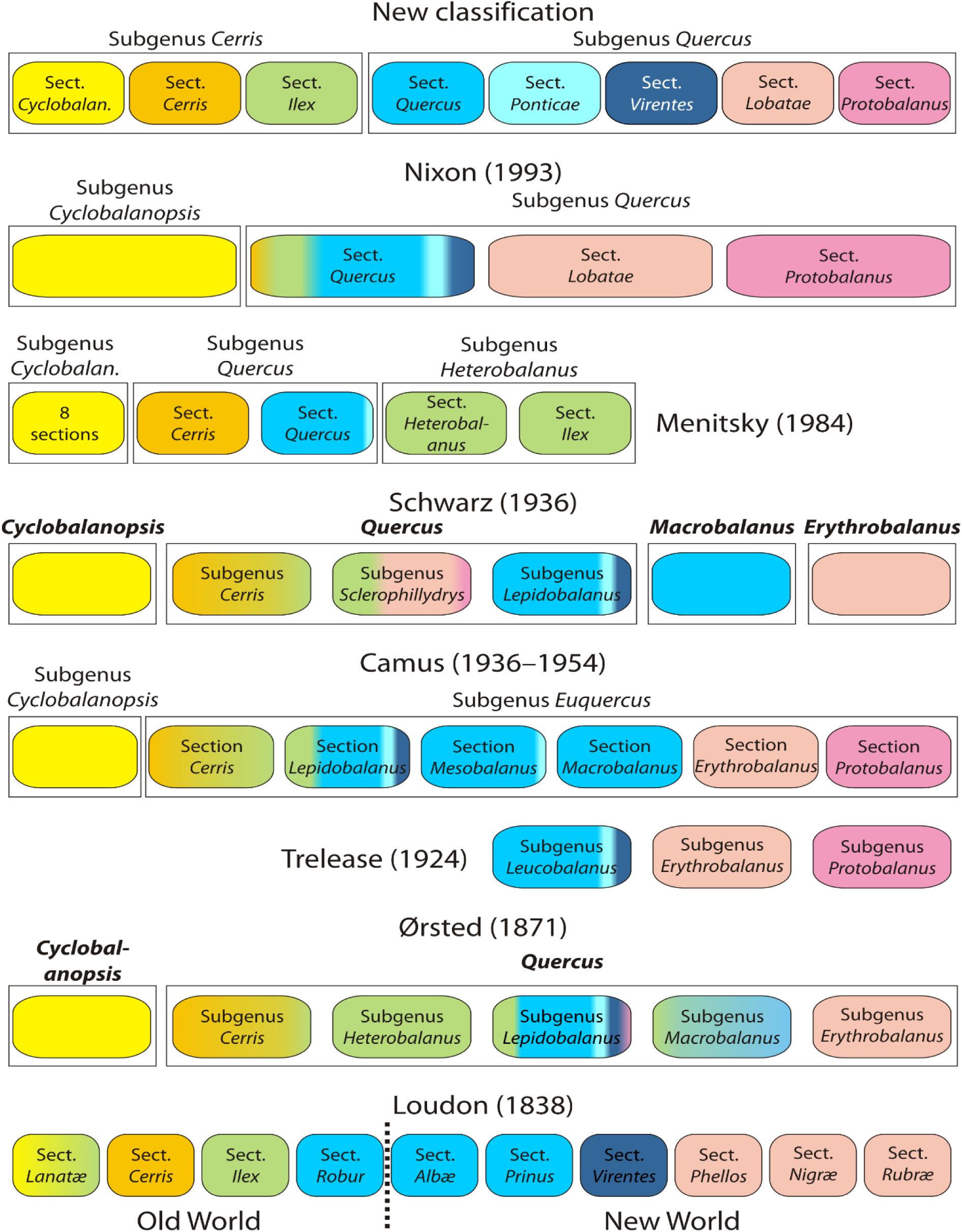
Classification schemes for *Quercus* from Loudon to Nixon (see Appendix). Colour coding denotes the actual systematic affiliation of species included in each taxon: of the ‘Old World’ or ‘mid-latitude clade’ section *Cyclobalanopsis* (cycle-cup oaks, yellow), section *Cerris* (Cerris oaks; orange), and section *Ilex* (Ilex oaks; green); and of the ‘New World’ or ‘high-latitude clade’ section *Quercus* (white oaks s.str.; blue), sections *Virentes* (cyan) and *Ponticae* (dark blue), section *Protobalanus* (intermediate oaks; purple), and section *Lobatae* (red oaks; red). Colour gradients are proportional, i.e. reflect the proportion of species with different systematic affiliation included in each taxon. Note: Menitsky (1984) and Trelease (1924) only treated the Eurasian and American oaks, respectively, and provided classifications in (nearly) full agreement with current phylogenies.

Ørsted (1871) can be credited for recognising an important Asian group of oaks hardly known at the time of Loudon and originally associated with *Cyclobalanus* (= *Lithocarpus*): the cycle-cup oaks of subtropical and tropical East Asia, which Ørsted considered distinct from *Quercus* as genus *Cyclobalanopsis*, within his subtribe Quercinae (**Fig. 1**). This concept was adopted by later researchers (e.g. Camus 1936–1938; Nixon 1993; as subgenera) and is still used for the Flora of China (Huang et al. 1999; Flora of China 2016). Within the second genus of the Quercinae, *Quercus,* Ørsted recognised five subgenera with a total of 16 sections and about 184 species. His work is the first to treat oaks in a global context; Loudon, and later Camus, Trelease, and Menitsky, treated the Nearctic and Palearctic-Indomalayan taxa independently.

In the early 20^th^ century, two competing classification concepts emerged, which were henceforth used by researchers (partly until today). The central/eastern European tradition followed in principle the classification system of Schwarz (1936), whereas the western/southern European tradition relied on the monographic work of Camus (1936–1938, 1938–1939, 1952–1954). A decade earlier, Trelease (1924) provided a comprehensive treatment of the North American oaks listing about 371 species (nearly half of them new) in 138 series and three subgenera/sections (**Fig. 1**): *Leucobalanus* (white oaks), *Erythrobalanus* (red oaks), and *Protobalanus* (intermediate oaks). Thus, he established the still valid tripartition of the genus in North America (sections *Quercus, Lobatae, Protobalanus*; (Jensen 1997; Manos 1997; Nixon and Muller 1997). Camus and Schwarz (partly) followed Trelease regarding the classification of North American oaks, but disagreed with respect to the oaks of Eurasia and North Africa, specifically on how to classify the North American oaks in relation to their Eurasian counterparts. Camus followed Ørsted’s general scheme, but recognised a single genus *Quercus* with the two subgenera *Cyclobalanopsis* and *Quercus.* She downgraded Ørsted’s subgenera in *Quercus* to sections (**Fig. 1**). Schwarz (1936) also followed in principle the concepts of Ørsted, but raised Ørsted’s categories, erecting a two-tribe system (Cyclobalanopsideae, Querceae) with two genera each (*Cyclobalanopsis + Erythrobalanus, Macrobalanus + Quercus*). A novelty in the system of Schwarz was the subgenus *Sclerophyllodrys* (**Fig. 1**), in which he accommodated many sclerophyllous oaks of Eurasia, Trelease’s subgenus *Protobalanus* (including an Asian series *Spathulatae*), and six evergreen series of Trelease’s subgenus *Erythrobalanus*. Another major difference relative to Camus was that Schwarz adopted Ørsted’s global concept by grouping North American and Eurasian white oaks in the same sections (*Dascia, Gallifera, Prinus, Roburoides*).

The most recent monographic work towards a new classification of oaks was the one of Menitsky (1984, translated into English in 2005) dealing with Asian oaks (**Fig. 1**). Except for a single species (*Q. suber*), Menitsky placed all Ilex oaks in subgenus *Heterobalanus*, while Cerris oaks (except for *Q. suber*) formed one of the two sections in subgenus *Quercus* (the other section included the white oaks). Menitsky’s account is the only morphology-based system that correctly identified the natural groups of Eurasian oaks confirmed later by palynological and molecular data. In the same way, Trelease’s sections of North American oaks also have been confirmed as natural groups.

The latest and currently most widely used (e.g. Govaerts and Frodin 1998; see also www.wikipedia.org and www.internationaloaksociety.org) classification is by Nixon (1993), published as a review. Nixon adopted the concept of Camus but merged her sections *Cerris,* which comprised Cerris and Ilex oaks, and *Euquercus,* comprising the remaining Ilex oaks and the white oaks, into a single section *Quercus.* According to this latest modification of Ørsted’s more than 150 years old scheme, the genus *Quercus* is divided into two subgenera, the cycle-cup oaks (*Cyclobalanopsis*) and all remaining oaks (*Quercus*). Subgenus *Quercus* includes two natural sections, one comprising the red oaks (sect. *Lobatae*) and one comprising the intermediate oaks (sect. *Protobalanus*), and a heterogeneous, artificial, northern hemispheric section *Quercus* including all white oaks, Cerris and Ilex oaks (**Fig. 1**).

### Change in criteria for classification

There are two major causes for the differences in the traditional, morphology-based classifications of oaks: 1) the weighing of morphological characters, 2) the geographic regions considered. Convergent morphological evolution is a common phenomenon in the genus *Quercus* and the Fagaceae in general (Oh and Manos 2008; Kremer et al. 2012). For instance, Loudon’s (1838) descriptions for the distantly related sections *Ilex* (Eurasian *Q. ilex* and relatives) and *Virentes* (North American *Q. virens* Ait. [= *Q. virginiana* Miller], a white oak relative) are essentially identical. For similar reasons, Ørsted (1871) included a section *Ilex* in his subgenus *Lepidobalanus* (white oaks in a broad sense), while expanding this section to include evergreen North American white oaks (the sect./subsect. *Virentes* of Loudon, Trelease, Camus, etc.) On the other hand, the Himalayan Ilex oak *Q. lanata* was included in Ørsted’s section *Prinus* of North American white oaks. The assumption that leaf texture can be used to assign species to higher taxonomic groups on a global scale supports Schwarz’ largely artificial subgenera (and genera to some degree). Using the descriptions by Trelease, the Eurasian Ilex oaks would still fall in his subgenus *Protobalanus*, and the same is true for the descriptions in Nixon (1993) and the Flora of North America (Manos 1997).

Nixon’s concept of a section *Quercus* including all white, Cerris and Ilex oaks primarily relies on the basal position of aborted ovules in these groups. Already de Candolle (1862b) noted this feature as being variable in different oak species, and Camus (1936–1938, p. 40f) emphasised that this trait is stable not only within a species, but also characterises groups of species (but see general descriptions in Menitsky 1984). Nixon also adopted Camus’ concept of subgenus *Cyclobalanopsis* (aborted ovules always apical; but see general description provided by Huang et al. 1999). Apical abortive ovules, on the other hand, are found in most but not all subsections of sect. *Erythrobalanus* (the red oaks) and in the castanoid genera.

Therefore, Nixon suggested that basal abortive ovules are a synapomorphy of his sect. *Quercus*. Subsequent work has shown that the position of aborted ovules in the mature seeds of *Quercus* is the result of different developmental processes and less stable than originally assumed (Borgardt and Pigg 1999; Borgardt and Nixon 2003; Deng et al. 2008; **Table 1**). The only two classification schemes that recognised the same groups later recovered in molecular studies are those by Trelease (1924) and Menitsky (1984). Notably, these monographs were restricted to North American and Eurasian oaks, respectively. Therefore, they did not run the risk of creating artificial groups including morphologically similar but unrelated Old World and New World species.

**Table 1:** Different contributions of placenta and funiculus to the position of aborted ovules in mature seeds of *Quercus*. Information compiled from Borgardt and Pigg (1999), Borgardt and Nixon (2003), Deng (2007), Deng et al. (2008), and complemented by Min Deng (unpublished data).

| Section(s)<br>Feature | *Quercus*,<br>*Ponticae*,<br>*Virentes* | *Lobatae* | *Protobalanus* | *Cyclobalanopsis* | *Cerris* | *Ilex* |
| --- | --- | --- | --- | --- | --- | --- |
| Position of aborted ovules | Basal | Apical<br>Type I | Apical, basal, or lateral | Apical<br>Types I, III | Apical, basal, or lateral<br>Type II | Basal or lateral<br>Types II, III |
| Placenta | Sessile | Elongated | ? | Elongated | Sessile (compressed) | Sessile or elongated |
| Funiculus | Sessile | Sessile | ? | Sessile or elongated | Sessile or elongated | Sessile or elongated |

Notes: **Type I**: apical/lateral aborted ovules by elongated placenta, **Type II**: by elongated funiculus, **Type III**: both elongated placenta and funiculus. Other Fagaceae (*Castanea*, *Castanopsis*, *Lithocarpus*, *Trigonobalanus*) have Types I & III aborted ovules. All other Fagaceae have apical aborted ovules.

### Changing from morphology to molecules

The first molecular phylogeny of *Quercus* including a comprehensive oak sample is the one of Manos et al. (2001) based on sequences of the nuclear ITS region and plastid RFLP data. While Manos et al.’s molecular phylogeny included only a limited sample of Old World species, it challenged the traditional views of Ørsted until Nixon. Instead, the intermediate and white oaks grouped with the red oaks, forming the ‘New World Clade’, but not with the Cerris and Ilex oaks. The latter formed an ‘Old World Clade’ that later would be shown to include the cycle-cup oaks (Manos et al. 2008). While the red oaks and cycle-cup oaks were resolved in well-supported and distinct clades within their respective subtrees, the situation appeared more complex for Camus’ section *Cerris* (including a few Ilex oaks) and the white oaks (Manos et al. 2001). The lack of unambiguous support may be one reason, why morphologists and oak systematists did not readily implement the new evidence (e.g. Borgardt and Nixon 2003; le Hardÿ de Beaulieu and Lamant 2010; see also www.internationaloaksociety.org). The other reason is probably that the two new clades lacked compelling, unifying morphological traits.

Plastid gene regions commonly used in plant phylogenetics turned out to be less useful for inferring infrageneric and inter- to intraspecific relationships in oaks. This is mainly because the plastid genealogy is largely decoupled from taxonomy and substantially affected by geography (e.g. Neophytou et al. 2010; Neophytou et al. 2011; Simeone et al. 2016; Pham et al. 2017). Using genus- to family-level plastid data sets, even when combined with nuclear data, oaks are consistently recognised as a diphyletic group. This is best illustrated in Manos et al. (2008): one moderately supported main clade comprises the ‘New World Clade’ of oaks and *Notholithocarpus*, a monotypic Fagaceae genus of western North America; the other major clade comprises the Eurasian Fagaceae *Castanea* and *Castanopsis*, and the ‘Old World Clade’ of *Quercus*. The phenomenon is also seen in broadly sampled plastid data sets and can produce highly artificial molecular phylogenies (e.g. Xiang et al. 2014; Xing et al. 2014) as discussed in Grímsson et al. (2016). Nevertheless, all currently available plastid data reject the traditional subdivision into two subgenera *Cyclobalanopsis* and *Quercus*: the overall signal (e.g. Manos et al. 2008) is in line with the ‘New World/Old World Clade’ concept introduced by Oh & Manos (2008).

In view of the problems encountered with plastid sequence data, oak molecular phylogenetics concentrated on nuclear-encoded sequence regions. Nine years after the study by Manos et al. (1999), the first ITS phylogeny was confirmed and supplemented by data from a single-copy nuclear gene region, the *Crabs Claw* (*CRC*) gene (Oh and Manos 2008). Denk & Grimm (2010) provided an updated Fagaceae ITS tree including more than 900 individual sequences of oaks (including ca. 600 newly generated for western Eurasian species taking into account substantial intra-individual variation). Their data on the 5S intergenic spacer (over 900 sequences), a multicopy nuclear rDNA gene region not linked with the ITS region, supported three groups of western Eurasian oaks as originally conceived by Menitsky (1984). Hubert et al. (2014) compiled new data from six single-copy nuclear gene regions and combined the new data with ITS consensus sequences (based on Denk and Grimm 2010) and *CRC* sequence data (Oh and Manos 2008). Most recently, Hipp et al. (2015) showed a tree based on a large, nuclear reduced representation next-generation sequencing (RADseq) data set. All these data sets and analyses support the recognition of two, reciprocally monophyletic groups of oaks (**Fig. 2**) that can be formalised as two subgenera with eight phylogenetic lineages (Hubert et al. 2014; Hipp et al. 2015), accepted here as sections that match the morphological groups originally perceived by Trelease (1924) and Menitsky (1984):

**Figure 2:**
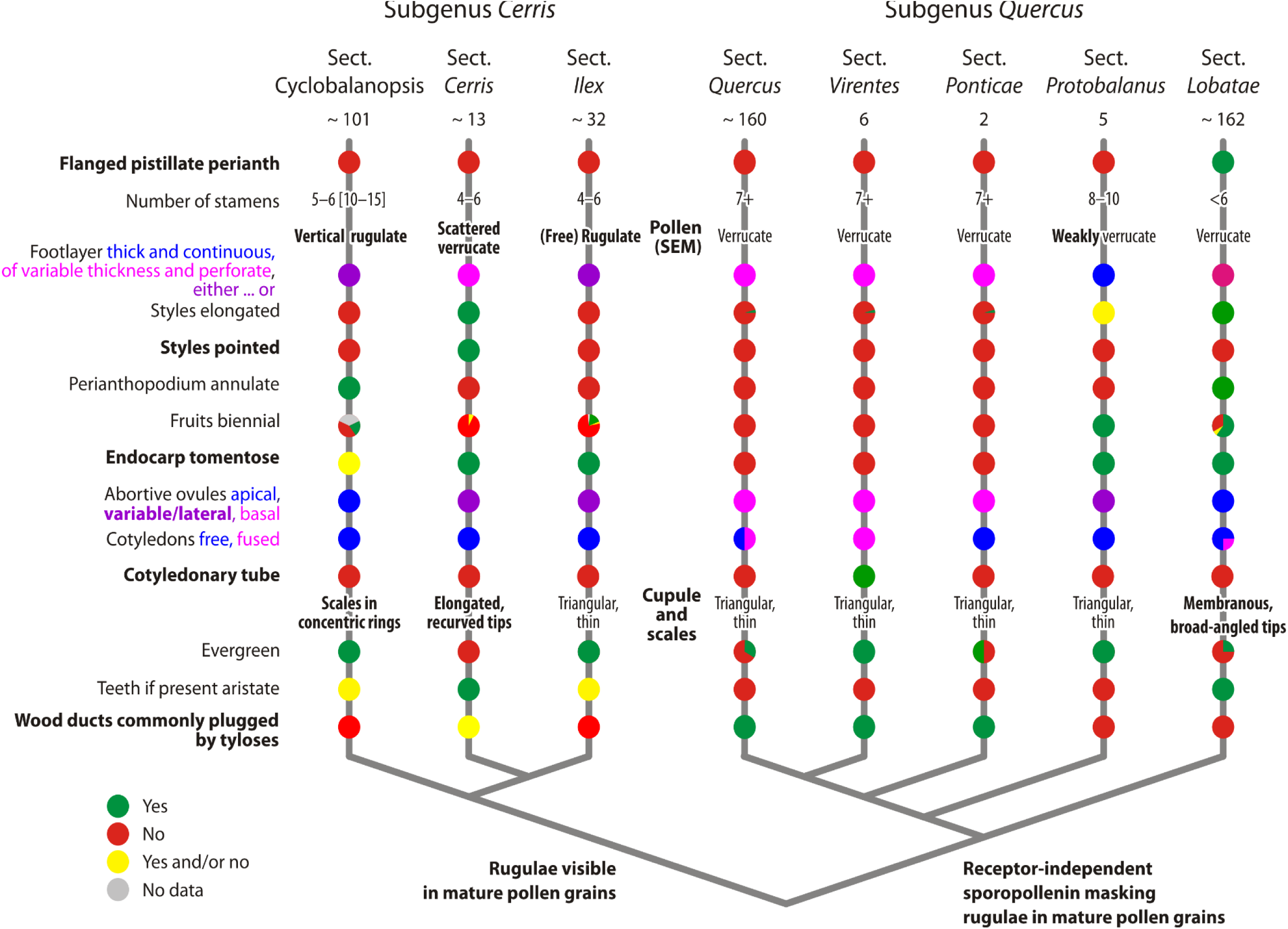
Revised sectional classification of oaks and diagnostic characters of lineages. The basic phylogenetic relationships of the six infrageneric groups of oaks are shown, formalised here as sections in two monophyletic subgenera, subgenus Cerris (‘Old World’ or ‘mid-latitude clade’) and subgenus Quercus (‘New World’ or ‘high-latitude clade’). Section-specific traits in bold; subgenus-diagnostic traits indicated at the respective branches of the schematic phylogenetic tree (Hubert et al. 2014; Hipp et al. 2015). Most traits are shared by more than one section of oaks including non-sister-lineages (normal font); they evolved convergently or are potentially plesiomorphic traits. Some are variable within a section as indicated by (semi-)proportional pie charts. Nonetheless, each section can be diagnosed by unique, unambiguous character suites. Note: ‘yes’ (green) and ‘no’ (red) refers to whether the mentioned trait is observed or not in members of the section, but should not be generally viewed as derived or ancestral.

- **Subgenus *Quercus***, the ‘New World clade’ (Manos et al. 2001) or ‘high-latitude clade’ (Grímsson et al. 2015; Simeone et al. 2016), including
  - the North American intermediate oaks, **section *Protobalanus*** (= Trelease’s subgenus of the same name);
  - the western Eurasian-western North American disjunct **section *Ponticae***;
  - the North American “southern live oaks”, **section** ***Virentes***;
  - all white oaks from North America (= Trelease’s subgenus *Leucobalanus*) and Eurasia (= Menitsky’s section *Quercus*), **section *Quercus***; and
  - the North American red oaks, **section *Lobatae*** (= Trelease’s subgenus *Erythrobalanus*).
- **Subgenus *Cerris,*** the exclusively Eurasian ‘Old World clade’ (or ‘mid-latitude clade’), including
  - the cycle-cup oaks of East Asia (including Malesia), **section *Cyclobalanopsis*** (former [sub]genus *Cyclobalanopsis* of Ørsted, Camus, Schwarz, Menitsky, and Nixon);
  - the Ilex oaks, **section *Ilex*** (= Menitsky’s subgenus *Heterobalanus* minus *Q. suber*); and
  - the Cerris oaks, **section *Cerris*** (= Menitsky’s section *Cerris* minus *Q. alnifolia*).

## Part B. Revised subgeneric and sectional classification of oaks

The following information for diagnostic morphological characters for the recognised groups of oaks is based mostly on information provided in Trelease (1924), Camus (1936–1938, 1938–1939, 1952–1954), Schwarz (1936, 1937), Menitsky (1984), le Hardÿ de Beaulieu & Lamant (2010), and the Floras of China (Huang et al. 1999) and North America (Flora of North America Editorial Commitee 1997). Information on pollen morphology is from Rowley et al. (1979), Solomon (1983a, 1983b), Rowley & Claugher (1991), Rowley (1996), Rowley & Gabarayeva (2004), Denk & Grimm (2009), Makino et al. (2009), and Denk & Tekleva (2014). Updated information on the position of aborted ovules and the relative contributions of placenta and funiculus to it is from Borgardt & Nixon (2003), Deng et al. (2008), and Min Deng (unpublished data).

If no reference is provided, most monographers (Trelease 1924; Schwarz 1936; Camus 1936–1938; Schwarz 1937; Camus 1938–1939, 1952–1954; Menitsky 1984) agreed on a particular character. Trelease (1924) emphasised the importance of wood characters for delimitation of major groups of North American oaks. According to Trelease (1924), Menitsky (1984), and Akkemik and Yaman (2012) the type of wood porosity and presence or absence of tyloses plugging vessels of early-wood are clade-specific to some degree.

Group-specific traits are highlighted by italics (see also **Fig. 2**).

### Genus *Quercus*

1753, Sp. Pl., 1: 994.

Lectotype: *Quercus robur* L. (selected by Britton and Brown, Ill. Fl. N. U.S. ed. 2. 1: 616, 7 Jun 1913; confirmed by Green, in Sprague, Nom. Prop. Brit. Bot.: 189, Aug 1929)

**Trees** 20–30(–55) m high, or **shrubs**; monoecious, evergreen or deciduous; **propagating** from seeds (saplings) or, occasionally, vegetative propagation (ramets); **bark** smooth or deeply furrowed or scaly or papery, corky in some species; **wood** ring-porous or (semi) diffuse-porous, tyloses common in vessels of early-wood or rarely present; **terminal buds** spherical to ovoid, terete or angled, all scales imbricate; **leaves** spirally arranged, stipules deciduous and inconspicuous or sometimes retained until the end of the vegetative period; **lamina** chartaceous or coriaceous, lobed or unlobed, margin entire, dentate or dentate with bristle-like extensions; **primary venation** pinnate; **secondary venation** eucamptodromous, brochidodromous, craspedodromous, semicraspedodromous, or mixed; intersecondary veins present or absent; **inflorescences** unisexual in axils of leaves or bud scales, usually clustered at base of new growth; **staminate inflorescences** lax, racemose to spicate; **pistillate inflorescence** usually stiff, a simple spike, with terminal cupule and sometimes one to several sessile, lateral cupules; **staminate flowers** subsessile, in dichasial clusters of 1–3(–7) (section *Cyclobalanopsis*) or solitary; sepals connate, stamens (2–)6(–15), pistillodes reduced and replaced by a tuft of silky hairs; **pollen** monad, medium-sized or small (size categories according Hesse et al. 2009), 3-colp(or)ate, shape prolate, outline in polar view trilobate or rounded, in equatorial view elliptic to oval, tectate, columellate; pollen ornamentation (micro)rugulate, (micro) rugulate-perforate, or (micro)verrucate, (micro)verrucate-perforate; foot layer discontinuous or continuous, of even or uneven thickness; **pistillate flower** one per cupule, with 1–2 subtending bracts, sepals connate, (3–)6 (–9) lobed, either situated directly on the tip of the ovary or on the perianthopodium (stylopodium); carpels and styles 3–6, occasionally with staminodes, styles with a broad stigmatic surface on adaxial suture of style (less prominent in section *Cyclobalanopsis*); **ovules** pendent, anatropous or semi-anatropous; **position of aborted ovules** apical, basal, or lateral depending on whether or not the placenta and/or funiculus are secondarily elongated; **fruit** a one-seeded nut (acorn) with a proximal scar, fruit maturation annual or biennial, nut one per cup, round in cross-section, not winged, cotyledons free or fused; **endocarp** glabrous or tomentose; **cup** covering at least base of nut, with lamellate rings or scaly; **scales** imbricate and flattened or tuberculate, not or weakly to markedly reflexed; **chromosome number X** = 12. Around 400 species mostly in the Northern Hemisphere (**Fig. 3**).

**Figure 3:**
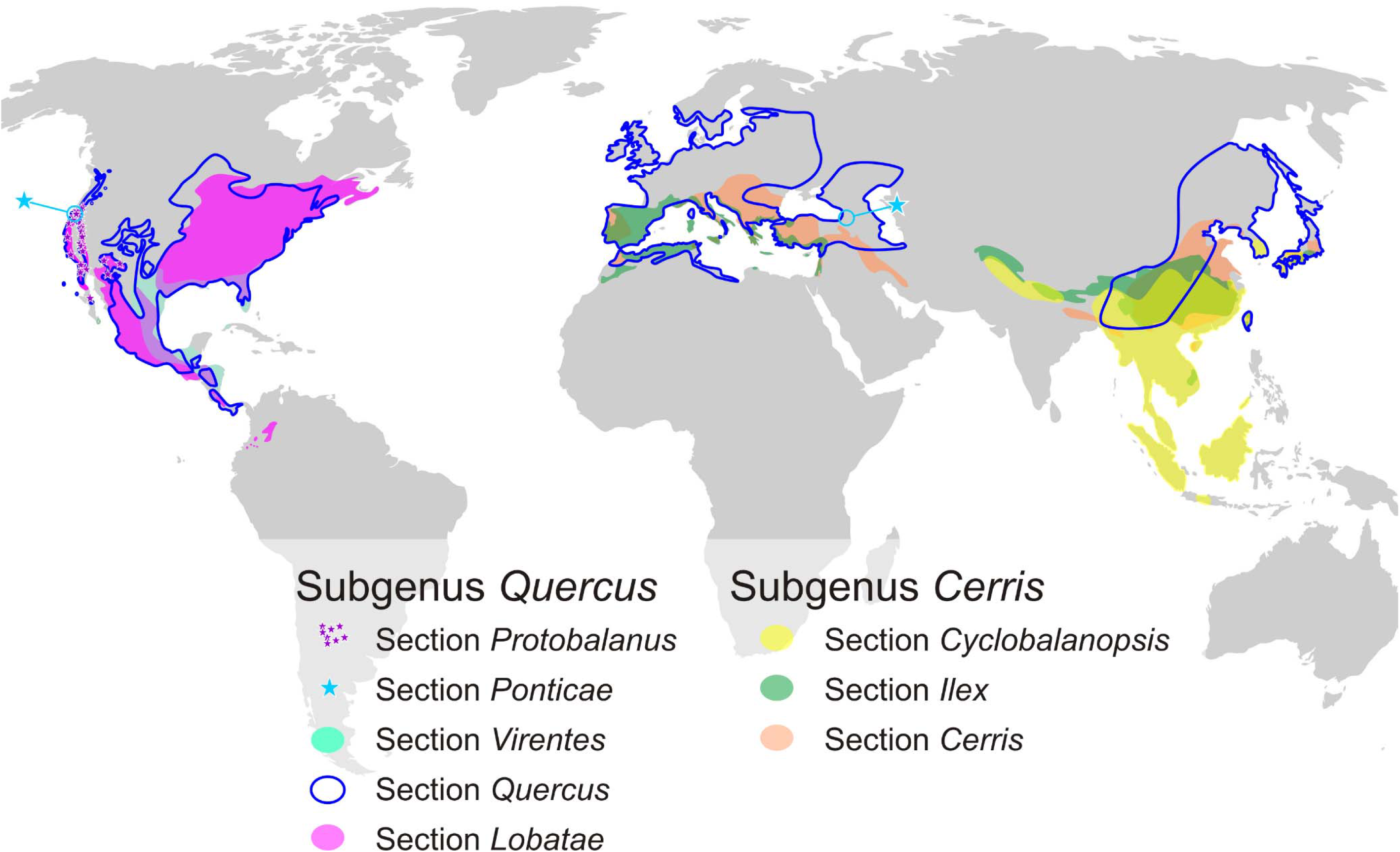
Geographic distribution of the eight sections of *Quercus*. Distribution data from Browicz and Zieliński (1982), Menitsky (1984), Costa Tenorio et al. (2001), Deng (2007), Fang et al. (2009), and Manos (2016)

#### 1. Subgenus *Quercus*

*Receptor-independent sporopollenin masking rugulae in mature pollen grains* (Rowley and Claugher 1991; Rowley 1996).

##### 1.1. Section *Protobalanus* (intermediate oaks)

*Quercus* section *Protobalanus* (Trelease) Schwarz, Notizbl. Bot. Gart. Berlin-Dahlem, 13/116: 21 (1936)

*Quercus* subgenus *Protobalanus* Trelease, in Standley, Contr. US Natl. Herb. 23:176 (1922). – *Quercus* section *Protobalanus* (Trelease) Camus, Les Chênes, 1: 157 (1938). – *Quercus* section *Protobalanus* (Trelease) Schwarz, Notizbl. Bot. Gart. Berlin-Dahlem, 13/116: 21 (1936) p.p.

Type: *Quercus chrysolepis* Liebm. (Trelease, Proc. Natl. Acad. Sci. 2: 627, 1916; confirmed by Nixon, Ann. Sci. For. 50, suppl. 1: 32s, 1993)

Stamens 8–10, with apiculate apices (Trelease 1924); *pollen ornamentation weakly verrucate*, perforate (Denk and Grimm 2009); footlayer thick and continuous (Denk and Tekleva 2014); styles short to long, elliptic in cross-section; stigmata abruptly dilated; stigmatic surface extending adaxially along stylar suture (Trelease 1924; Manos 1997); fruit maturation biennial (Trelease 1924; Camus 1952–1954; Manos 1997); endocarp tomentose (Trelease 1924; Camus 1952–1954; Manos 1997); position of abortive ovules basal, lateral or apical, can be variable within a single plant (Manos 1997); cup scales triangular and fused at the base, thickened and compressed into rings, often tuberculate and obscured by glandular trichomes, with sharp angled tips; leaf dentitions spinose; wood diffuse porous, tyloses rarely present in vessels of early-wood (Trelease 1924).

Five species in southwestern North America and northwestern Mexico (Manos 1997).

##### 1.2. Section *Ponticae*

*Quercus* section *Ponticae* Stefanoff., Ann. Univ. Sofia, ser. 5, 8: 53 (1930)

*Quercus* ser. *Sadlerianae* Trelease, Oaks of America: 111 (1924). – *Quercus* subsect. *Ponticae* Menitsky (Stefanoff) A.Camus, Bull. Soc. Bot. Fr., 81: 815 (1934). – *Quercus* ser. *Ponticae* Schwarz, Notizbl. Bot. Gart. Berlin-Dahlem, 13/116: 11 (1936).

Lectotype (here designated): *Quercus pontica* K.Koch.

Shrubs or small trees, rhizomatous; number of stamens mostly 6 (Trelease 1924; Camus 1952–1954); pollen ornamentation verrucate (Denk and Grimm 2009); footlayer of variable thickness and perforate (Denk and Tekleva 2014); staminate catkins up to 10 cm long; styles short, fused or free, elliptic in cross-section; stigmata abruptly or gradually dilated (Schwarz 1936); fruit maturation annual; endocarp glabrous; position of abortive ovules basal; cotyledons free; cup scales slightly tuberculate with sharp angled apices, occasionally with attenuated tips (Trelease 1924; Gagnidze et al. 2014); leaves evergreen or deciduous, chestnut-like, stipules large, persistent or early shed, number of secondary veins 10–15(–25), dentate, teeth simple or compound (in *Q. pontica*), sharply mucronate or with thread-like, curved upwards extension; leaf buds large, *bud scales loosely attached* (Trelease 1924; Schwarz 1936; Menitsky 1984); wood ring porous or diffuse porous, large vessels commonly plugged by tyloses.

Two species in mountainous areas of north-eastern Turkey and western Georgia (Transcaucasia) and in western North America (northern-most California, southern-most Oregon; Trelease 1924; Menitsky 1984; Gagnidze et al. 2014) (**Fig. 3**).

##### 1.3. Section *Virentes*

*Quercus* section *Virentes* Loudon, Arbor. Frut. Brit., 3: 1918 (1838)

*Quercus* ser. *Virentes* Trelease, Oaks of America: 112 (1924)

Type: *Quercus virens* Aiton (= *Q. virginiana* Mill.)

Trees or rhizomatous shrubs; pollen ornamentation verrucate (Denk and Grimm 2009); footlayer of variable thickness and perforate (Denk and Tekleva 2014); styles short, fused or free, elliptic in cross-section; stigmata abruptly or gradually dilated (Schwarz 1936); fruit maturation annual; cup scales narrowly triangular, free or fused at the base, thinly keeled and barely tuberculate with sharp angled apices; leaves evergreen or subevergreen (Trelease 1924; Nixon and Muller 1997); wood diffuse porous, tyloses abundant in large vessels (Trelease 1924); *cotyledons fused* (de Candolle 1862a; Engelmann 1880); *germinating seed with elongated radicle/epicotyl forming a tube* (Nixon 2009); hypocotyl region produces a tuberous fusiform structure.

Ca. 7 species in south-eastern North America, Mexico, the West Indies (Cuba), and Central America (Muller 1961; Cavender-Bares et al. 2015) (**Fig. 3**).

##### 1.4. Section *Quercus* (white oaks s.str.)

*Quercus* section *Albae* Loudon, Arbor. Frut. Brit., 3: 1730 (1838). – *Quercus* section *Prinus* Loudon, Arbor. Frut. Brit., 3: 1730 (1838). – *Quercus* section *Robur* Loudon, Arbor. Frut. Brit., 3: 1731 (1838). – *Quercus* section *Virentes* Loudon, Arbor. Frut. Brit., 3: 1730 (1838). – *Quercus* section *Gallifera* Spach, Hist. Nat. Veg., 11:170 (1842). – *Quercus* section *Eulepidobalanus* Oerst., Vidensk. Meddel. Naturhist. Foren. Kjøbenhavn 1866, 28: 65 (1866–1867) p.p. – *Quercus* section *Macrocarpae* Oerst., Vidensk. Meddel. Naturhist. Foren. Kjøbenhavn 1866, 28: 68 (1866–1867). – *Quercus* section *Diversipilosae* C.K.Schneid., Handb. Laubholzk., 1: 208 (1906). – *Quercus* section *Dentatae* C.K.Schneid., Handb. Laubholzk., 1: 209 (1906). – *Quercus* section *Mesobalanus* A.Camus, Bull. Soc. Bot. Fr., 81: 815 (1934). – *Quercus* section *Roburoides* Schwarz, Notizbl. Bot. Gart. Berlin-Dahlem 13: 10 (1936). – *Quercus* section *Robur* Schwarz, Notizbl. Bot. Gart. Berlin-Dahlem 13: 12 (1936). – *Quercus* section *Dascia* (Kotschy) Schwarz, Notizbl. Bot. Gart. Berlin-Dahlem 13: 14 (1936).

Stamens ≥ 7 (Trelease 1924; Camus 1936–1938, 1938–1939); pollen ornamentation verrucate (Denk and Grimm 2009); footlayer of variable thickness and perforate (Denk and Tekleva 2014); styles short, fused or free, elliptic in cross-section; stigmata abruptly or gradually dilated; stigmatic surface extending adaxially along stylar suture (all authors; best illustrated in Schwarz 1936, fig. 1); fruit maturation annual; endocarp glabrous or nearly so; cotyledons free or fused; position of abortive ovules basal (de Candolle 1862b; confirmed/accepted by later authors), placenta and funiculus sessile; cup scales triangular, free or fused at the base, thickened, keeled and often tuberculate with sharp angled apices, occasionally with attenuated tips; leaf dentitions typically without bristle-like, aristate tips; wood ring porous, large vessels in (early-)wood commonly plugged by tyloses (Trelease 1924; see Akkemik and Yaman 2012).

Ca. 150 species in North America, Mexico, Central America, western Eurasia, East Asia, and North Africa (Nixon and Muller 1997).

##### 1.5. Section *Lobatae* (red oaks)

*Quercus* section *Lobatae* Loudon, Hort. Brit., 385 (1830)

*Quercus* section *Integrifoliae* Loudon, Hort. Brit., 384 (1830) p.p. – *Quercus* section *Mucronatae* Loudon, Hort. Brit., 385 (1830) p.p. – *Quercus* section *Rubrae* Loudon, Arbor. Frut. Brit., 3: 1877 (1838; see also Loudon 1839). – *Quercus* section *Nigrae* Loudon, Arbor. Frut. Brit., 3: 1890 (1838; see also Loudon 1839). – *Quercus* section *Phellos* Loudon, Arbor. Frut. Brit., 3: 1894 (1838; see also Loudon 1839). – *Quercus* section *Erythrobalanus* Spach, Hist. veg. Phan., 11:160 (1842). – *Quercus* subgenus *Erythrobalanus* (Spach) Oerst., Vidensk. Meddel. Naturhist. Foren. Kjøbenhavn, 28: 70 (1866–1867). – *Erythrobalanus* (Spach) O.Schwarz (as genus), Notizbl. Bot. Gart. Berlin-Dahlem 13: 8 (1936).

Lectotype: *Quercus aquatica* (Lam.) Walter (= *Q. nigra* L.) (Nixon, Ann. Sci. For. 50, suppl.1: 30s, 1993)

*Pistillate perianth forming a characteristic flange* (Schwarz 1936, fig. 1; Nixon 1993; Jensen 1997); number of stamens ≤ 6 (Trelease 1924; Camus 1952–1954); pollen ornamentation verrucate (Denk and Grimm 2009); footlayer of variable thickness and perforate (Denk and Tekleva 2014); styles elongated, linear, outcurved, elliptic in cross-section; stigmata slightly dilated, spatulate to oblong; stigmatic surface extending adaxially along stylar suture (Trelease 1924); perianthopodium conical, often annulate (Trelease 1924); fruit maturation biennial, rarely annual (le Hardÿ de Beaulieu and Lamant 2010); endocarp tomentose; cotyledons free or sometimes basally fused; position of abortive ovules apical or rarely lateral to basal (de Candolle 1862b; Trelease 1924), placenta sessile or elongated, funiculus sessile; *cupule fused with peduncle forming a ‘connective piece’* (compare Denk and Meller 2001) for *Fagus*), connective piece covered with small scales similar to those on the cupule; *cup scales* triangular and free, mostly thin, membranous and smooth *with broadly angled tips*; *leaf teeth and lobes typically with bristle-like extensions*, teeth reduced to bristles in entire or nearly entire leaves; wood ring-porous or semi ring-porous, late-wood markedly porous, tyloses in vessels of early-wood rarely present (Trelease 1924).

Ca. 120 species in North America, Mexico, Central America, and Colombia in South America (Jensen 1997).

#### 2. Subgenus *Cerris*

*Quercus* subgenus *Cerris* Oerst., Vidensk. Meddel. Naturhist. Foren. Kjøbenhavn 1866, 28: 77 (1866–1867)

Lectotype (here designated): *Quercus cerris* L.

*Rugulae visible in mature pollen grains or weakly masked* (Solomon 1983a, b; Denk and Grimm 2009; Makino et al. 2009; Denk and Tekleva 2014).

##### 2.1. Section *Cyclobalanopsis*

*Quercus* sect. *Cyclobalanopsis* (Oerst.) Benth. & Hook. f., Gen. Plant. 3, 408 (1880)

*Cyclobalanopsis* Oerst. (as genus), Vidensk. Meddel. Naturhist. Foren. Kjøbenhavn 1866, 28: 77 (1866–1867), nom. conserv. – *Quercus* sect. *Cyclobalanopsis* (Oerst.) Benth. & Hook. f., Gen. Plant. 3, 408 (1880). – *Quercus* subgenus *Cyclobalanopsis* (Oerst.) Schneider, Ill. Handb. Laubholzk. 1, 210 (1906).

Type: *Quercus velutina* Lindl. ex Wall., non Lam. (vide Farr and Zijlstra 2017)

Staminate flowers in groups of 1–3(–7) along inflorescence axis (Menitsky 1984; Nixon 1993); stamens 5–6 (Huang et al. 1999) to 10–15 (Ohwi 1965); *pollen ornamentation vertical-rugulate* (Denk and Grimm 2009); footlayer thick and continuous or of variable thickness and perforate (Denk and Tekleva 2014); styles short to very short (< 3mm to < 1mm), elliptic in cross-section; stigmata dilated, subcapitate; stigmatic surface not forming a prominent stigmatic groove (Camus 1936–1938; Menitsky 1984; Nixon 1993; Huang et al. 1999); *perianthopodium annulate with 3-5 distinct rings* (Schwarz 1936); fruit maturation annual or biennial (Camus 1936–1938; le Hardÿ de Beaulieu and Lamant 2010); endocarp tomentose ore rarely glabrous (Camus 1936–1938); cotyledons free; position of abortive ovules apical (Camus 1936–1938; Menitsky 1984) [note: according to Huang et al. (1999) the position is variable, but no details are provided in the species descriptions], placenta elongated reaching the apical part of the seed, where vascular bundles enter the seed and the aborted ovules, funiculus sessile or with short petiole; *cupule with concentric lamellae*; leaves evergreen; leaf dentitions with bristle-like extensions or not; wood diffuse porous, tyloses very rarely present in vessels of early-wood (Menitsky 1984).

Ca. 90 species in tropical and subtropical Asia including the southern Himalayas (Huang et al. 1999).

##### 2.2. Section *Ilex*

*Quercus* section *Ilex* Loudon, Arbor. Frut. Brit., 3: 1899 (1838)

*Quercus* subgenus *Heterobalanus* Oerst., Vidensk. Meddel. Naturhist. Foren. Kjøbenhavn 1866, 28: 69 (1866–1867). – *Quercus* subgenus *Heterobalanus* (Oerst.) Menitsky, Duby Azii, 89 (1984) [Oaks of Asia, 133, (2005)]. – *Quercus* section *Heterobalanus* (Oerst.) Menitsky, Duby Azii, 89 (1984) [Oaks of Asia, 134, (2005)]. – *Quercus* subsection *Ilex* (Loudon) Guerke sensu Menitsky, Duby Azii, 97 (1984) [Oaks of Asia, 151, (2005)].

Type: *Quercus ilex* L.

Stamens 4 to 6 (Schwarz 1937); *pollen ornamentation rugulate* (Denk and Grimm 2009); footlayer thick and continuous or of variable thickness and perforate (Denk and Tekleva 2014); styles medium-long, apically gradually dilated, recurved, v-shaped in diameter; stigmata slightly subulate; stigmatic surface extending adaxially along stylar suture (Schwarz 1937; Menitsky 1984); fruit maturation annual or biennial (Camus 1936–1938, 1938–1939; Menitsky 1984; le Hardÿ de Beaulieu and Lamant 2010) [note that observations by Menitsky partly differ from those of Camus and le Hardÿ de Beaulieu and Lamant]; endocarp tomentose (Schwarz 1936; Camus 1936–1938; Schwarz 1937; Camus 1938–1939; Menitsky 1984); cotyledons free; position of the abortive ovules basal or lateral, placenta and funiculus sessile or elongated; cup scales triangular, free or fused at the base, mostly thin, membranous, often keeled and tuberculate with sharp angled apices, occasionally with slightly raised tips or narrowly triangular, well-articulated, thickened with elongated recurved tips (as in *Q. alnifolia, Q. baronii, Q. coccifera, Q. dolicholepis*); leaves evergreen, dentitions spinose or with bristle-like extensions; wood diffuse porous, tyloses rarely present in vessels of early-wood (Menitsky 1984).

Ca. 35 species in Eurasia and North Africa (Menitsky 1984; Denk and Grimm 2010; Deng et al. 2017).

##### 2.3. Section *Cerris*

*Quercus* section *Cerris* Dumort., Florula Belgica: 15 (1829).

*Quercus* section *Cerris* Loudon, Arbor. Frut. Brit., 3: 1730 (1838). – *Quercus* section *Eucerris* Oerst., Vidensk. Meddel. Naturhist. Foren. Kjøbenhavn 1866, 28: 75, nom. illeg. (1867). – *Quercus* section *Erythrobalanopsis* Oerst., Vidensk. Meddel. Naturhist. Foren. Kjøbenhavn 1866, 28: 76 (1867). – *Quercus* section *Castaneifolia* O.Schwarz, Feddes Repert., 33: 322 (1934). – *Quercus* section *Vallonea* O.Schwarz, Feddes Repert., 33: 322 (1934). – *Quercus* section *Aegilops* (Reichenb.) O.Schwarz, Notizbl. Bot. Gart. Berlin-Dahlem, 13/116: 19 (1936).

Type: *Quercus cerris* L.

Number of stamens 4–6; *pollen ornamentation scattered verrucate* (Denk and Grimm 2009); footlayer of variable thickness and perforate (Denk and Tekleva 2014); *styles* elongated, outcurved*, pointed*, v-shaped in diameter; stigmatic area linear; stigmatic surface extending adaxially along stylar suture; fruit maturation biennial, variable only in *Q. suber* (Camus 1936–1938; le Hardÿ de Beaulieu and Lamant 2010); endocarp tomentose (Camus 1936–1938); cotyledons free; position of abortive ovules basal, lateral or apical (de Candolle 1862b; Camus 1936–1938; Schwarz 1937; Menitsky 1984), placenta sessile, funiculus sessile or elongated; *cup scales* narrowly triangular, well-articulated, thickened and keeled *with elongated, well-developed recurved tips*; leaf dentitions typically with bristle-like extensions; wood (semi-)ring-porous, tyloses in vessels of early-wood present but not common (Trelease 1924; Akkemik and Yaman 2012).

Ca. 10–12 species in Eurasia and North Africa (Menitsky 1984).

### Infrasectional classification: the big challenge

The main challenge for oak systematics in the coming years will be a meaningful classification below the sectional rank. Nuclear-phylogenomic data (Hipp et al. 2015) within *Quercus* recover subclades that occur in well-defined biogeographic regions (e.g. western North America, eastern North America, western Eurasia, East Asia etc.), a sorting not so clear from traditional sequence data. Only very few New World-Old World disjuncts are recognised (e.g. section *Ponticae*; McVay et al. 2017). Ongoing phylogenomic work (Hipp et al. 2014; Hipp et al. 2015; McVay et al. 2017; Hipp et al. submitted) is beginning to reveal structure within the sections *Lobatae* and *Quercus* that corresponds to regional diversity within the Americas. Preliminary phylogenetic analyses suggested that the early evolutionary branches of *Lobatae* include many of the lobed-leaf species groups of North America (Hipp et al. 2015). The first branch, however, comprises the seven Californian taxa (*Agrifoliae* sensu Trelease), followed by various groups containing mostly temperate species that sort out into well-defined subclades. For section *Quercus*, analyses suggest some uncertainty at the base of the clade, specifically regarding the position of the Eurasian subclade. Previous morphology-based treatments of the Eurasian white oaks (‘roburoids’) suggested close affinities to certain eastern North American species, like *Q. montana* Willd. (series *Prinoideae* of Trelease), based on a similar (e.g. ‘prinoid’) leaf morphology (Axelrod 1983). In the most recent time calibrated tree (Hipp et al. submitted) the roburoids are nested within the white oak (s.str.) clade and diverged at around 25–30 Ma (Oligocene) from a North American, fully temperate clade including the type species of the section *Quercus* (*Q. alba*) and morphologically similar species such as *Q. montana* (see also Pearse and Hipp 2009; McVay et al. 2017). Dispersed pollen provide evidence for the presence of the white oak s.l. lineage (pollen types found today in sections *Quercus* and *Ponticae* but not *Virentes*) in the middle Eocene of western Greenland and the Baltic amber region of northern Europe (Grímsson et al. 2015; Grímsson et al. 2016; Eva-Maria Sadowski, unpublished data). Hence, the early radiation of this lineage involved the North Atlantic land bridge. Pollen and leaf fossil evidence from Eocene and Oligocene strata in East Asia further indicate migration from North America via the Bering land bridge. The expansion of this East Asian branch of white oaks (early roburoids) gave rise to the (modern) western Eurasian roburoids. This is in some agreement with the latest dated tree proposing a crown age of ca. 15–20 Ma for the roburoids. By that time, the final radiation within the North American white oaks (s.str.; section *Quercus*) might have been completed (Hipp et al. submitted).

Lack of resolution using traditional sequence data, e.g. identical ITS variants found in North American and Eurasian white oaks, and the relative young inferred root (stem) and crown ages of the roburoids can be explained by recent episodic migration from North America to Europe across the North Atlantic land bridge in addition to probably (very) large population sizes of temperate (white) oaks. Ancient hybridisation between roburoids and section *Ponticae* has been identified as one possible source of potentially misleading phylogenetic data (McVay et al. 2017).

However, the current, partly preliminary results also indicate that (sub)sections/series recognised in the monographs of Trelease, Camus, and Menitsky do not always correspond to groups identified when using molecular sequence data. In some cases, the molecular-defined groups may seem counterintuitive. For example, the western Eurasian Ilex oaks *Quercus alnifolia, Q. aucheri, Q. coccifera* and *Q. ilex* are resolved as a monophyletic group (Denk and Grimm 2010; Hipp et al. 2015), but were placed into two subgenera and three sections by Schwarz (1936), and two sections and three subsections by Camus (1936–1938, 1938–1939) and Menitsky (1984) due to conspicuous differences in indumentum, leaf margin, and cup scales. Similar mismatches between traditional classification and DNA-based groups are encountered for all large infrageneric groups and will pose a major challenge when searching for morphological criteria to subdivide sections within oaks. For example, while it has long been noticed that characters of the indumentum of the abaxial leaf surface provide valuable information for species delimitation (Manos 1993; Nixon 2002; Tschan and Denk 2012; Deng et al. 2015, 2017), these characters appear to have evolved convergently in related and unrelated groups (Tschan and Denk 2012; Deng et al. 2017).

## Part C. Fossil record of the eight sections

Section *Protobalanus—*In addition to pollen of section *Lobatae*, Grímsson et al. (2015) found dispersed pollen similar to pollen of section *Protobalanus* in middle Eocene deposits of western Greenland. Unambiguous leaf fossils and dispersed pollen of section *Protobalanus* are known from the latest Eocene-earliest Oligocene Florissant Formation (ca. 34 Ma; Bouchal et al. 2014). The section had a western and northern North American distribution during the Paleogene and became restricted to western North America during the Neogene.

Section *Ponticae*—Miocene and Pliocene leaf fossils assigned to *Quercus pontica miocenica* Kubát have traditionally been compared with the extant *Q. pontica* (e.g. Andreánszky 1959; Gagnidze et al. 2014). Presently, these fossils are included within the fossil-species *Q. gigas* Göppert, a fossil representative of section *Cerris* (Walther and Zastawniak 1991; Kvaček et al. 2002). These leaf remains and early Cenozoic fossils from Arctic regions (see Grímsson et al. 2016) are superficially similar to leaves of section *Ponticae* but lack the characteristic dentition of *Q. pontica*. Hence, there is currently no reliable fossil record of this section because chestnut-like foliage might have evolved in parallel in various modern and extinct lineages of Fagaceae or represents the primitive state within genus *Quercus* or subgenus *Quercus*.

Section *Virentes*—For similar reasons as outlined above for section *Ponticae* there is no reliable fossil record of this modern section.

Section *Quercus—*Leaf fossils from middle Eocene deposits of Axel-Heiberg Island, Canadian Arctic (ca. 45 Ma; McIver and Basinger 1999) most likely belong to section *Quercus*. The leaf fossils are strikingly similar to the extant East Asian *Quercus aliena* Blume var. *acutiserrata* Maxim. ex Wenzig. From the roughly coeval Baltic amber of northern Europe, Crepet (1989) described *in situ* pollen of male flowers, which may represent section *Quercus*. Fagaceous remains from the Baltic amber are currently revised by Eva-Maria Sadowski, Göttingen, and may represent more than one section of *Quercus*. The leaf fossil-taxon *Quercus kraskinensis* Pavlyutkin from the early Oligocene of the Primorskii Region (Pavlyutkin 2015) is similar to the material from Axel-Heiberg Island and most likely belongs to section *Quercus*. The Paleogene radiation of the white oak lineage might have involved both the Bering and the North Atlantic land bridges. Lobed oaks of section *Quercus* are also known from the Oligocene of Central Asia and Northeast Asia (e.g. Krishtofovich et al. 1956; Tanai and Uemura 1994), and North America (Bouchal et al. 2014). Hence, the section had a scattered distribution during the Paleogene that included low, mid, and high latitudes. In the Neogene, section *Quercus* was widespread across the entire Northern Hemisphere (Borgardt and Pigg 1999).

Section *Lobatae—*Oldest fossils that can securely be assigned to section *Lobatae* are dispersed pollen from the middle Eocene of western Greenland (latest Lutetian to earliest Bartonian, 42–40 Ma). The pollen shows a sculpturing found today only in members of section *Lobatae* (Grímsson et al. 2015). A further record of entire-margined, lanceolate foliage with preserved epidermal structures from middle Eocene (48–38 Ma) deposits of Central Europe – described as *Quercus subhercynica* (Kvaček and Walther 1989) and originally assigned to section *Lobatae –* was subsequently transferred to the extinct genus *Castaneophyllum*, a fossil-genus for which morphological affinities to certain castaneoid genera have been established using leaf epidermal characteristics (Kvaček and Walther 2010 [2012]). For North America, Daghlian and Crepet (1983) described cups and acorns associated with lobate leaves from the Oligocene (Rupelian, ca. 30 Ma) Catahoula Formation in Texas (*Quercus oligocenensis*). Markedly similar leaf records from the early Oligocene of northeast Asia, *Quercus sichotensis* Ablaev et Gorovoj, *Q. ussuriensis* Krysh., *Q. arsenjevii* Ablaev et Gorovoj, and *Q. kodairae* Huzioka assigned to section *Cerris* (Tanai and Uemura 1994; Pavlyutkin 2015) clearly also belong to section *Lobatae*. Hence, the section had a mid to high latitude northern hemispheric distribution during the Paleogene; during the Neogene it was present in Europe (Jähnichen 1966; Kovar-Eder and Meller 2003) and North America.

Section *Cerris—*The earliest unambiguous record of section *Cerris, Quercus gracilis* (Pavlyutkin) Pavlyutkin, comes from early Oligocene leaf fossils from the Russian Far East (Pavlyutkin et al. 2014). In western Eurasia, earliest evidence of section *Cerris* comes from dispersed pollen from late Oligocene/ early Miocene (ca. 23 Ma) deposits of Central Europe (Kmenta 2011). The section has a rich Neogene fossil record in Eurasia (e.g. Mai 1995; Song et al. 2000; Yabe 2008).

Section *Ilex*—The Paleogene record of Ilex oaks is so far limited to dispersed pollen from East Asia (Hainan Island, China, Changchang Formation, Lutetian-Bartonian, ca. 40 Ma; Hofmann 2010; Spicer et al. 2014) and Central Europe (Germany, Rupelian, ca. 33 Ma; Denk et al. 2012). From the Changchang Formation, Spicer et al. (2014) also reported leaf morphotypes (OTUs 68 and 71) that most likely belong to section *Ilex*. There is also leaf fossil evidence of section *Ilex* from 26 Ma strata in Tibet (Zhou Zhekun, personal communication). Section *Ilex* has a rich fossil record in Neogene deposits across Eurasia (e.g. Denk et al. 2017).

Section *Cyclobalanopsis—Quercus paleocarpa* (Manchester 1994) cupules and nuts from the Eocene (Lutetian, ca. 48 Ma) of western North America are possibly the oldest fossils belonging to section *Cyclobalanopsis*, but without preserved stigmas the assignment of these fruits remains ambiguous. Additionally, fossilised fruits of *Cyclobalanopsis nathoi* from the middle Eocene of Japan (Huzioka and Takahashi 1973) may belong to section *Cyclobalanopsis* based on the shape of nuts. Hofmann (2010) reported dispersed pollen grains from the middle Eocene of Hainan Island, China (Changchang Formation, Lutetian-Bartonian, 48–38 Ma; Spicer et al. 2014). From the Changchang Formation, Spicer et al. (2014) also reported leaf morphotypes (OTUs 61, 62 [partly], 63, 67) that most likely belong to section *Cyclobalanopsis*. From the Oligocene of south-western China, several fossil-species based on leaf impressions have been assigned to cycle cup oaks (Writing Group of Cenozoic Plants of China [WGCPC] 1978). Evidence for assignment to section *Cyclobalanopsis* is based on the number, arrangement, and course of secondary veins, the dentition, and the attenuate leaf apex (e.g. *Q. parachampionii* Cheu and Liu; *Q. paraschottkyana* Wang and Liu). This section has a Paleogene record in mid to low latitude East Asia and western North America, while it is restricted to Asia during the Neogene (e.g. Jia et al. 2015). No reliable records are known from Europe.

## Conclusion and outlook

Recent molecular phylogenetic studies consistently suggest two major clades within oaks, one comprising three Old World groups (sections *Cyclobalanopsis, Ilex*, and *Cerris*), the other comprising three New World groups (sections *Protobalanus, Virentes*, and *Lobatae*,) and two northern hemispheric groups (sections *Ponticae* and *Quercus*). This is in contrast to the established view that *Cyclobalanopsis* oaks are sister to the remainder of the genus *Quercus*. The reason for this conflict is that morphological characters evolved convergently in all major groups of oaks and even outside oaks in other Fagaceae (e.g. concentric cupula rings). Important conserved morphological and diagnostic characters are pollen sculpturing and ultrastructure (**Fig. 2**).

Based on the new molecular and morphological evidence the infrageneric classification of *Quercus* is revised. A major challenge for future studies will be the molecular and morphological circumscription of infrasectional groups and their biogeographic and ecological characterisation. In this context, comparative morphological investigations of the seed ontogeny will be important to document the distribution of type I, II and III developmental pathways of aborted ovule positions (**Table 1**) across all sections. Some characters that have been described mainly on the basis of herbarium material, such as the annual or biennial mode of maturation, need to be reinvestigated in the field. In (fully) evergreen species of sections *Ilex, Protobalanus*, and *Cyclobalanopsis,* the fruiting twigs do not produce new growth in the second year after pollination (pseudo-annual maturation sensu Nixon 1997) and therefore may erroneously be interpreted as annual. In a number of high-mountain species of section *Ilex*, maturation may take much longer than previously assumed, with time periods of up to three to four years between pollination and mature seeds (Min Deng, unpublished data). Considering the oak fossil record, it is noteworthy that Paleogene plant-bearing deposits from East Asia and the Far East have so far been understudied. Once recovered, this fossil record should contribute to a better understanding of the emergence of major groups within oaks.

## Acknowledgements

We thank John McNeill for valuable comments. This work was supported by the Swedish Research Council (VR, grant to TD). GWG acknowledges financial support by the AMS Wien.

## Appendix

The appendix is provided as a spread-sheet file (XLSX) including eight spread-sheets, uploaded as supplementary information, and containing the following information:

- an <Overview> of earlier and current systematic schemes for oaks (genera, subgenera, sections);
- characters used/reported by earlier systematicists (sheets <Loudon>, <Trelease>, <Camus>, <Schwarz>, <Menitsky>, <90s>) extracted from the original literature;
- a comprehensive <All species> list of formerly and currently accepted species of oaks, compiled from the cited oak monographs and complemented by further data sources.

A copy of the appendix, which may be subject to future updates, will be included in an online archive provided for download at www.palaeogrimm.org/data/Dnk17_Appendix.zip.

Feel free to contact the authors for suggestions regarding (or errors found) in the appendix.

